# Analyses of the complete genome sequence of 2,6-dichlorobenzamide (BAM) degrader *Aminobacter* sp. MSH1 suggests a polyploid chromosome, phylogenetic reassignment, and functions of (un)stable plasmids

**DOI:** 10.1101/2021.02.25.432982

**Authors:** Tue Kjærgaard Nielsen, Benjamin Horemans, Cedric Lood, Jeroen T’Syen, Vera van Noort, Rob Lavigne, Lea Ellegaard-Jensen, Ole Hylling, Jens Aamand, Dirk Springael, Lars Hestbjerg Hansen

## Abstract

*Aminobacter* sp. MSH1 (CIP 110285) can use the pesticide dichlobenil and its transformation product, the recalcitrant groundwater micropollutant, 2,6-dichlorobenzamide (BAM) as sole source of carbon, nitrogen, and energy. The concentration of BAM in groundwater often exceeds the threshold limit for drinking water, resulting in the use of additional treatment in drinking water treatment plants (DWTPs) or closure of the affected abstraction wells. Biological treatment with MSH1 is considered a potential sustainable alternative to remediate BAM-contamination in drinking water production. Combining Illumina and Nanopore sequencing, we here present the complete genome of MSH1, which was determined independently in two different laboratories. Unexpectedly, divergences were observed between the two genomes, i.e. one of them lacked four plasmids compared to the other. Besides the circular chromosome and the two previously described plasmids involved in BAM catabolism pBAM1 (41 kb) and pBAM2 (54 kb), we observe that the genome of MSH1 contains two megaplasmids pUSP1 (367 kb) and pUSP2 (366 kb) and three smaller plasmids pUSP3 (97 kb), pUSP4 (64 kb), and pUSP5 (32 kb). The MSH1 substrain from KU Leuven showed a reduced genome lacking plasmids pUSP2 and the three smaller plasmids and was designated substrain MK1, whereas the variant with all plasmids was designated as substrain DK1. Results of a plasmid stability experiment, indicate that strain MSH1 may have a polyploid chromosome when growing in R2B medium with more chromosomes than plasmids per cell. Based on phylogenetic analyses, strain MSH1 is reassigned as *Aminobacter niigataensis* MSH1.

**Importance:** The complete genomes of the two MSH1 substrains, DK1 and MK1, provide further insight into this already well-studied organism with bioremediation potential. The varying plasmid contents in the two substrains suggest that some of the plasmids are unstable, although this is not supported by the herein described plasmid stability experiment. Instead, results suggest that MSH1 is polyploid with respect to its chromosome, at least under some growth conditions. As the essential BAM-degradation genes are found on some of these plasmids, stable inheritance is essential for continuous removal of BAM. Finally, *Aminobacter* sp. MSH1 is reassigned as *Aminobacter niigataensis* MSH1, based on phylogenetic evidence.

## Introduction

The occurrence of organic micropollutants in different water compartments threatens both ecosystem functioning as well as future drinking water supplies (1). Organic micropollutants are organic chemicals with complex and highly variable structures, and they have in common that they occur in the environment at trace concentrations (in the µg – ng/L range). Organic micropollutants often have unknown ecotoxicological and/or human health effects. They include a multitude of compounds such as pharmaceuticals, pesticides, ingredients of household products and additives of personal care products. In the European Union, the threshold limit for pesticides and relevant transformation products in drinking water is set at 0.1 µg/L (2). This threshold is frequently exceeded and forces drinking water treatment plants to invest in expensive physicochemical treatment technologies or to close groundwater extraction wells (3). The use of pollutant degrading bacteria in bioaugmentation strategies to remove micropollutants, such as pesticides, from drinking water, is presented as a solution (3, 4). The groundwater micropollutant 2,6-dichlorobenzamide (BAM), a transformation product of the herbicide dichlobenil, frequently occurs in groundwater in Europe, often exceeding the treshold concentration (5). *Aminobacter* sp. MSH1 (CIP 110285) was enriched and isolated from dichlobenil treated soil sampled from the courtyard of a plant nursery in Denmark. The strain converts dichlobenil to BAM, which is further fully mineralized (6). Efforts to elucidate the catabolic pathway for BAM degradation in MSH1 revealed the involvement of two plasmids. The first step of BAM-mineralization involves the hydrolysis of BAM to 2,6-dichlorobenzoic acid (2,6-DCBA) by the amidase BbdA encoded on the 41 kb IncP1-β plasmid pBAM1 (7). Further catabolism of 2,6-DCBA to central metabolism intermediates involves enzymes encoded on the 54 kb *repABC* family plasmid pBAM2. The strain mineralizes BAM at trace concentrations (6) and invades biofilms of microbial communities of rapid sand filters used in DWTPs (8). Moreover, it was successfully used in bioaugmentation of rapid sand filters, both in lab scale and pilot scale biofilration systems, to remove BAM from (ground)water (8–11). On the other hand, long-term population persistence and catabolic activity in the sand filters were impeded, likely due to a combination of predation and wash out (11, 12), as well as to physiological and genetic changes. Reducing flow rate and improving inoculation strategy have demonstrated prolonged persistence and activity of MSH1 in bioaugmented sand filters (13). However, other studies indicate that MSH1 shows a starvation survival response, in the nutrient (especially carbon) limiting environment of DWTPs, leading to reduced specific BAM degrading activity (14). Moreover, a substantial loss of plasmid pBAM2 was observed upon prolonged transfer of MSH1 both in R2A medium and in C-limited minimal medium (15), indicating that the plasmid is not entirely stable. Moreover, mutants lacking the ability to convert BAM into 2,6-DCBA have been reported (7). Clearly, to come to full management of bioaugmentation using MSH1 in DWTP biofiltration units aiming at BAM removal, more knowledge is needed on the physiological as well as genetic adaptations of MSH1 when introduced into the corresponding oligotrophic environment. The elucidation of the full genome sequence is crucial in this.

The complete genome sequence presented in this study shows that MSH1 substrain DK1 has a single chromosome and seven plasmids, including the two previously described catabolic plasmids pBAM1 and pBAM2, while substrain MK1 lacks four of these plasmids. The relative sequence coverage of the plasmids compared to the chromosome suggested that there are either multiple copies of the chromosome per cell or that there are, on average, fewer than one copy of six out of the seven plasmids per cell. This was tested in a plasmid stability experiment with substrain DK1 where plasmids were found to be overall stable, with the exception of a single loss event of pUSP1. This supports the hypothesis that MSH1 might have a polyploid chromosome, at least under some growth conditions.

## Material and Methods

### Growth conditions, genomic DNA preparation and sequencing

The genome sequence of strain MSH1 was independently obtained in two different laboratories, i.e., the KU Leuven in Belgium (MK1) and the Aarhus University lab in Roskilde, Denmark (DK1). In both cases, *Aminobacter* sp. MSH1 was obtained from the strain collection of the laboratory that originally isolated the bacterium (6). Sequencing of substrain DK1 at the Roskilde lab was performed as follows. Directly derived from a cryostock obtained from the original lab of MSH1, two ml of a culture grown in R2B were used for extraction of high molecular weight (HMW) DNA using the MasterPure^TM^ DNA Purification Kit (Epicentre, Madison, WI, USA), using the kit’s protocol for cell samples. DNA was eluted in 35 µL 10 mM Tris-HCl (pH 7.5) with 50 mM NaCl. The purity and concentration of extracted DNA were measured with a NanoDrop 2000c and a Qubit® 2.0 fluorometer (Thermo Fisher Scientific, Walther, MA, USA), respectively. An Illumina Nextera XT library was prepared for paired-end sequencing on an Illumina NextSeq 500 with a Mid Output v2 kit (300 cycles) (Illumina Inc., San Diego, CA, USA). Paired-end reads (2×151 bp) were trimmed for contaminating adapter sequences and low quality bases (<Q20) at the ends of the reads were removed using Cutadapt (v1.8.3) (16). Paired-end reads that overlapped were merged with AdapterRemoval (v2.1.0) (17). For Oxford Nanopore sequencing, a library was prepared from the same DNA extract using the Rapid Sequencing kit (SQK-RAD004). This was loaded on an R9.4 flow cell and sequenced using MinKnow (v1.11.5) (Oxford Nanopore Technologies, Oxford, UK). Nanopore reads were basecalled with albacore (v2.1.10) without quality filtering of reads. Only reads longer than 5,000 bp were retained and sequencing adapters were trimmed using Porechop (v0.2.3). A hybrid genome assembly with Nanopore and Illumina reads was performed using Unicycler (v0.4.3) (18).

The Illumina sequencing of substrain MK1 in the KU Leuven lab was reported previously (7, 19). Briefly, genomic DNA was isolated from a culture grown on R2B using the Puregene Core kit A (Qiagen, Hilden, Germany), according to the manufacturer’s instructions, except that DNA precipitation was performed with ethanol. A library was constructed for paired-end sequencing using 500 bp inserts and sequencing was performed on the Illumina GAIIx platform. Generated read lengths were 90 bp. The Illumina reads were quality controlled using FastQC (20) (v0.11.6) and BBduk (21) (v36.47). This included trimming the reads with low scoring regions (Phred < 30), clipping adapters, and removing very short reads (length < 50). For Nanopore sequencing, total genomic DNA was extracted from a culture grown on R2B with 200 mg/L BAM using the DNeasy UltraClean Microbial Kit (Qiagen, Hilden, Germany). Afterwards, the genomic DNA was mechanically sheared using a Covaris g-Tube (Covaris Inc., MA, USA) to an average fragment length of 8 kb. The library for sequencing was prepared using the 1D ligation approach with native 1D barcoding (SQK-LSK109) and sequenced on a MinION R9.4 flow cell using the MinION sequencer (Oxford Nanopore Technologies, Oxford, UK). The Nanopore reads were basecalled with Albacore (v2.0.2), and the barcode sequences were trimmed using Porechop (v0.2.3). Hybrid assembly of genome was performed as reported above.

### Genome analyses

For both genomes, automatic gene annotation was done using Prokka (22) (v1.14.0). Separately from Prokka, proteins with transmembrane helices were identified using TMHMM v2.0 (23). Genes were assigned to COG functional categories using EggNOG-mapper v4.5.1 (24). Genome comparison was done using EDGAR (25). Metabolic pathways were explored using Pathway Tools (26) and RAST (27). Circularized views of chromosome and plasmids were made using Circos (28). MegaX (29) was used for protein alignment and tree building. Phylogenetic analysis for strain MSH1 was performed using a clustal-omega (30) multiple sequence alignment using 16S ribosomal RNA genes from the set of type strains available in the *Phyllobacteriaceae* family. The tree was inferred using PhyML (31) with a GTR substitution model and a calculation of branch support values (bootstrap value of 1,000). Whole-genome-based taxonomic classification was performed with *in silico* DNA:DNA hybridization using the Type Strain Genome Server (TYGS) (32). Furthermore, average nucleotide identity (ANI) values were calculated for MSH1 against all available *Aminobacter* genomes in NCBI (downloaded January 31, 2021), using FastANI (33) and plotted in R with the *pheatmap* package (34). Genomes of the two MSH1 substrains were compared using the Mauve genome alignment software (35). Plasmids were characterized with regards to relaxase genes and replicon families using MOB-suite (36).

### Plasmid (in)stability experiment

To test for plasmid stability, MSH1 cells from -80*°*C cryostock were streaked on R2A plates and DNA from 1 ml of the cryostock was extracted using MasterPure^TM^ DNA Purification Kit. After incubation at 22°C for 11 days, a single colony from the R2A plate was picked and resuspended in 105 µl phosphate-buffered saline (PBS). From this, 5 µl suspension was inoculated in 25 ml R2B for 72 hours. Whole genome sequencing was performed on the remaining 100 µl PBS suspension. After 72 hours of growth in R2B, 1 mL broth culture was sampled for DNA extraction, similarly to the DNA extracted for initial DNA sequencing (above), and 100 µL of dilution series 10^-5^-10^-8^ of the R2B culture were plated onto R2A and the plates incubated at 22°C. After 7 days of growth, DNA was extracted and sequenced, as described above, from 14 individual colonies (originating from a single cell) resuspended in 100 µL PBS. All sequencing was performed on an Illumina NextSeq 550 with a Mid Output v2 kit (300 cycles) using Nextera XT library preparations as described above.

Sequencing adapters and poor quality sequences were trimmed from paired end reads using Trimmomatic (v0.39) (37) with the options “ILLUMINACLIP:/usr/share/trimmomatic/NexteraPE-PE.fa:2:30:10 LEADING:3 TRAILING:3 SLIDINGWINDOW:4:15 MINLEN:36”. Trimmed and filtered reads from each replicate MSH1 sample were mapped with bwa (v0.7.17-r1198-dirty) (38) to the completely assembled MSH1 genome including plasmids pBAM1-2 and pUSP1-5. Sequencing coverage in 1,000 bp windows for all replicons per replicate sample was calculated with samtools (v1.9-166-g74718c2) (39) and bedtools (v2.28.0) (40). Coverage data for all replicons were divided by the mean coverage of the chromosome, in order to normalize replicon copy numbers relative to the chromosome. Normalized coverage of all replicons for all replicates were visualized with Circos (v0.69-6) (28).

### Data availability

The genome sequences of strain MSH1 substrains MK1/DK1 are available under the following GenBank accession numbers CP026265/CP028968 (chromosome), CP026268/CP028967 (pBAM1), CP026267/CP028966 (pBAM2), CP026266/CP028969 (pUSP1) and CP028970 (pUSP2), CP028971 (pUSP3), CP028972 (pUSP4) and CP028973 (pUSP5).

## Results and discussion

### Genome statistics

The MSH1 genome (based on substrain DK1) consists of a chromosome of 5,301,518 bp and seven plasmids. The genome contains two large plasmids pUSP1 of 367,423 bp and pUSP2 of 365,485 bp, three smaller plasmids pUSP3, pUSP4, and pUSP5 (respectively 97,029 bp, 64,122 bp, and 31,577 bp) and the two previously reported smaller catabolic plasmids pBAM1 and pBAM2 of 40,559 bp and 53,893 bp, respectively (Table 1). A total of 6,257 genes could be predicted of which six rRNAs, 53 tRNAs, and four ncRNAs. A total of 6,194 CDS were predicted including 190 pseudogenes (Table 2). Circular views of the chromosome and seven plasmids are shown in Figure 1 and 2. The KU Leuven variant, designated as substrain MK1, lacked one of the two larger plasmids, i.e. pUSP2, and the three smaller plasmids pUSP3, pUSP4, and pUSP5. Except for the discrepancy in plasmids, the shared genomes (chromosome, pUSP1, pBAM1, and pBAM2) of the two strains have an average nucleotide identity of 99.9925%. The BAM-catabolic genes were manually checked for mutations that could indicate differences in degradation potential. A single nucleotide change was noted in the *bbdb3* gene on pBAM2, encoding one of three subunits of a TRAP-type transport system potentially involved in the uptake of 2,6-DCBA (19). In this gene, a non-synonomous substitution has changed a glycine to an arginine in the resulting protein in MK1. Currently, it is not known if this change has an effect on the putative function of this tripartite transport system. Furthermore, differences were found in the region of plasmid pUSP1 containing an IS*30* family insertion sequence with 38 bp flanking, imperfect, inverted repeats (IRs). The repeats appear complete in DK1, but MK1 shows a deletion of 56 bp and 34 bp up- and downstream of the IS*30* transposase gene, including partial deletion of the IR at both ends, suggesting that the MK1 substrain has undergone further genetic changes. The deletions flanking the IS*30* element on pUSP1 in MK1 may have been caused by a possible intramolecular transposition event (41). However, this IS*30* element with deletion in the IRs in MK1 may still be functional, as the functional core region of IS*30* IRs are only part of the complete IR (42).

**Figure 1.**
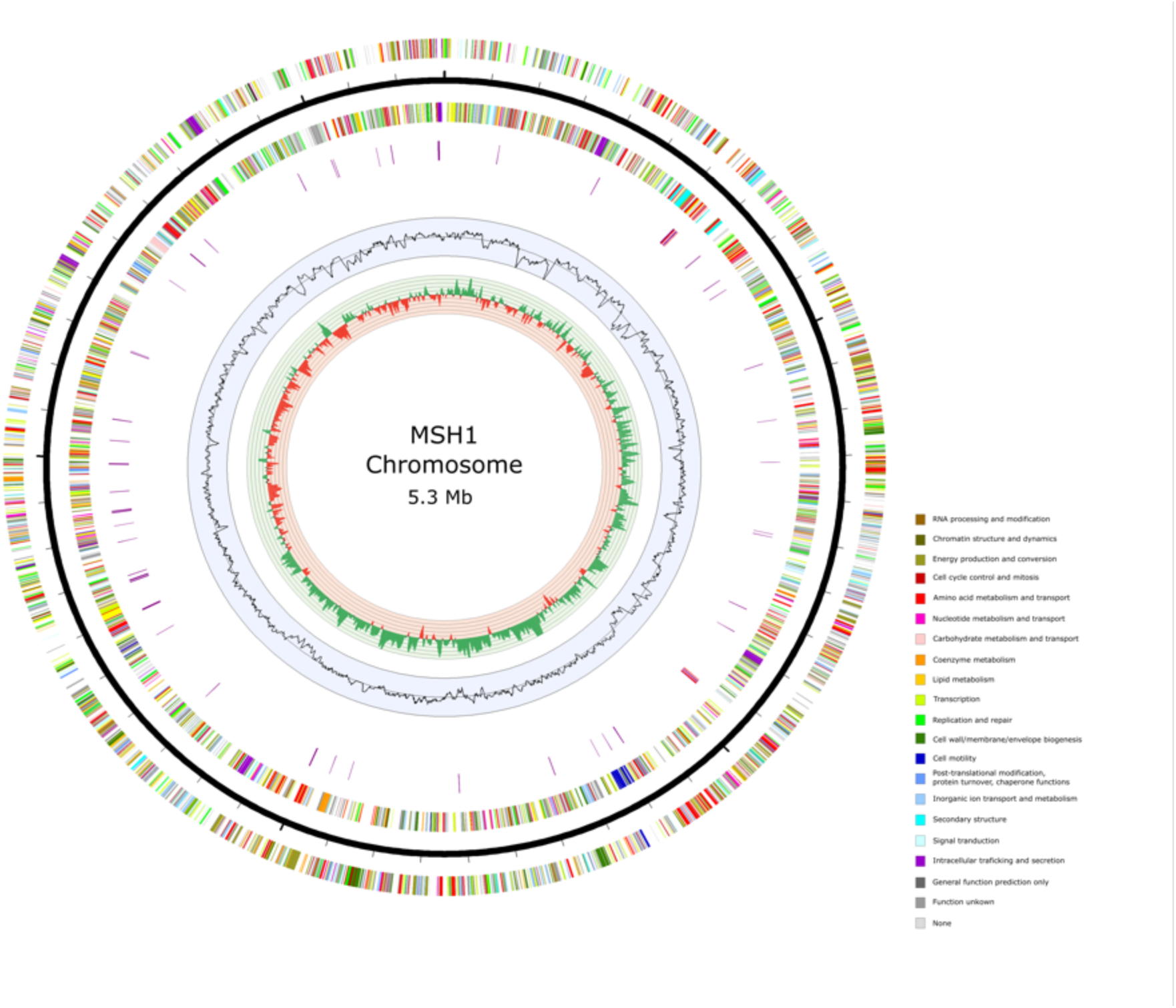
Circular view of the chromosome of *Aminobacter* sp. MSH1. From outer to inner circle: CDS on leading strand, scale (ticks: 100 kb), CDS on lagging strand, tRNA (purple) and rRNA (red) (only chromosome), GC plot and GC skew (>0: green, <0: red). CDS are colored according to COG functional categories determined with EggNOG mapper 4.5.1.

**Figure 2.**
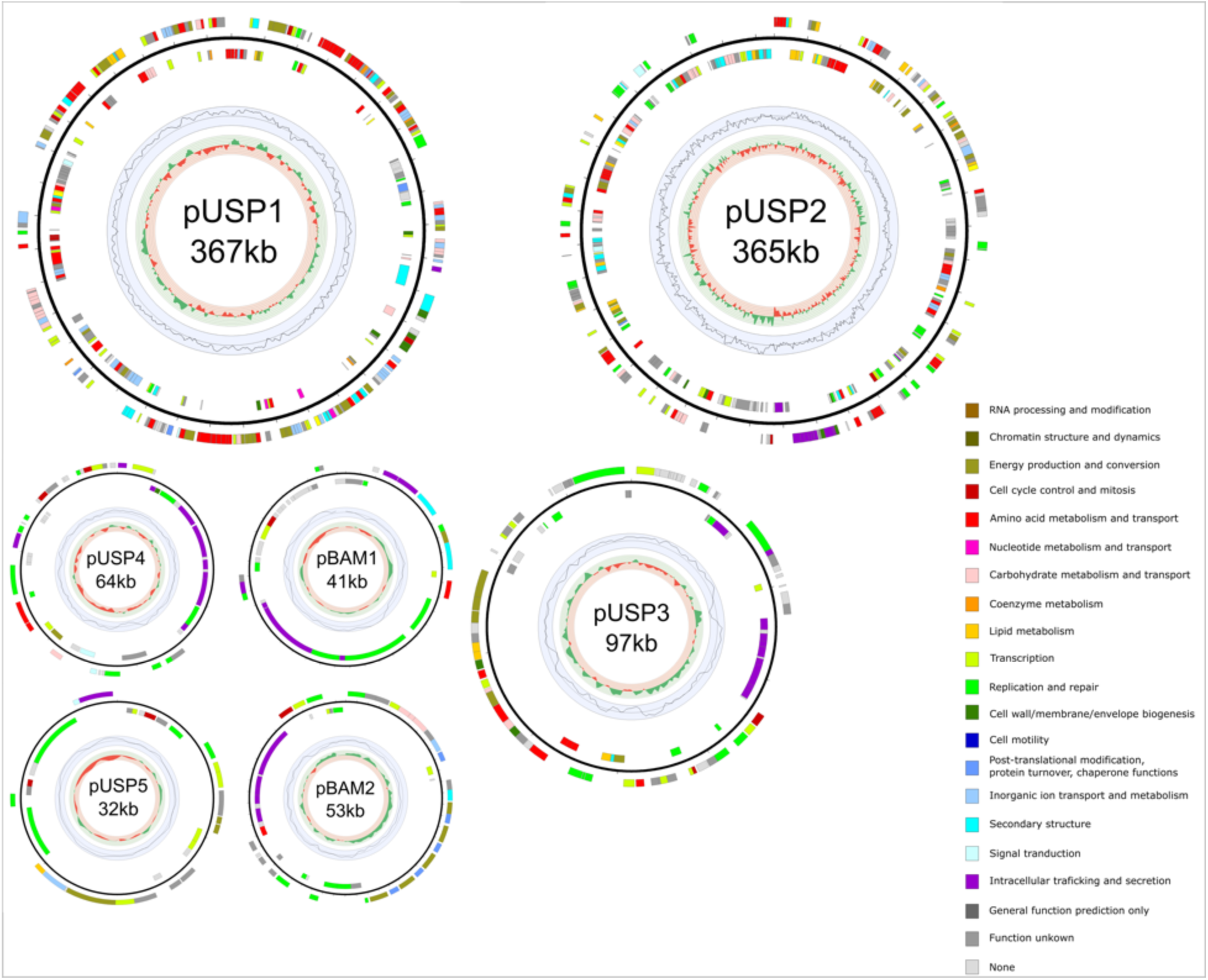
Circular view of the plasmids of the newly assigned *Aminobacter niigataensis* MSH1. From outer to inner circle: CDS on leading strand, scale (ticks: 100 kb), CDS on lagging strand, GC plot and GC skew (>0: green, <0: red). CDS are colored according to COG functional categories determined with EggNOG mapper 4.5.1. The KU Leuven substrain MK1 lacks plasmids pUSP2-5.

**Table 1.** Genome accession codes

| Label | Size (Mb) | GC (%) | Topology | INSDC identifier | RefSeq ID |
| --- | --- | --- | --- | --- | --- |
| Chromosome | 5.30 | 63.2 | Circular | CP028968.1(CP026265.1)* | NZ_CP028968.1(NZ_CP026265.1)* |
| Plasmid 1 pBAM1 | 0.04 | 64.4 | Circular | CP028967.1(CP026268.1)* | NZ_CP028967.1(NZ_CP026268.1)* |
| Plasmid 2 pBAM2 | 0.05 | 56.0 | Circular | CP028966.1(CP026267.1)* | NZ_CP028966.1(NZ_CP026267.1)* |
| Plasmid 3 pUSP1 | 0.37 | 63.1 | Circular | CP028969.1(CP026266.1)* | NZ_CP028969.1(NZ_CP026266.1)* |
| Plasmid 4 pUSP2 | 0.37 | 60.1 | Circular | CP028970.1 | NZ_CP028970.1 |
| Plasmid 5 pUSP3 | 0.10 | 60.5 | Circular | CP028971.1 | NZ_CP028971.1 |
| Plasmid 6 pUSP4 | 0.06 | 61.9 | Circular | CP028972.1 | NZ_CP028972.1 |
| Plasmid 7 pUSP5 | 0.03 | 62.9 | Circular | CP028973.1 | NZ_CP028973.1 |

\* INSDC identifier and RefSeqID of KU Leuven substrain MK1 submission in brackets

**Table 2.**
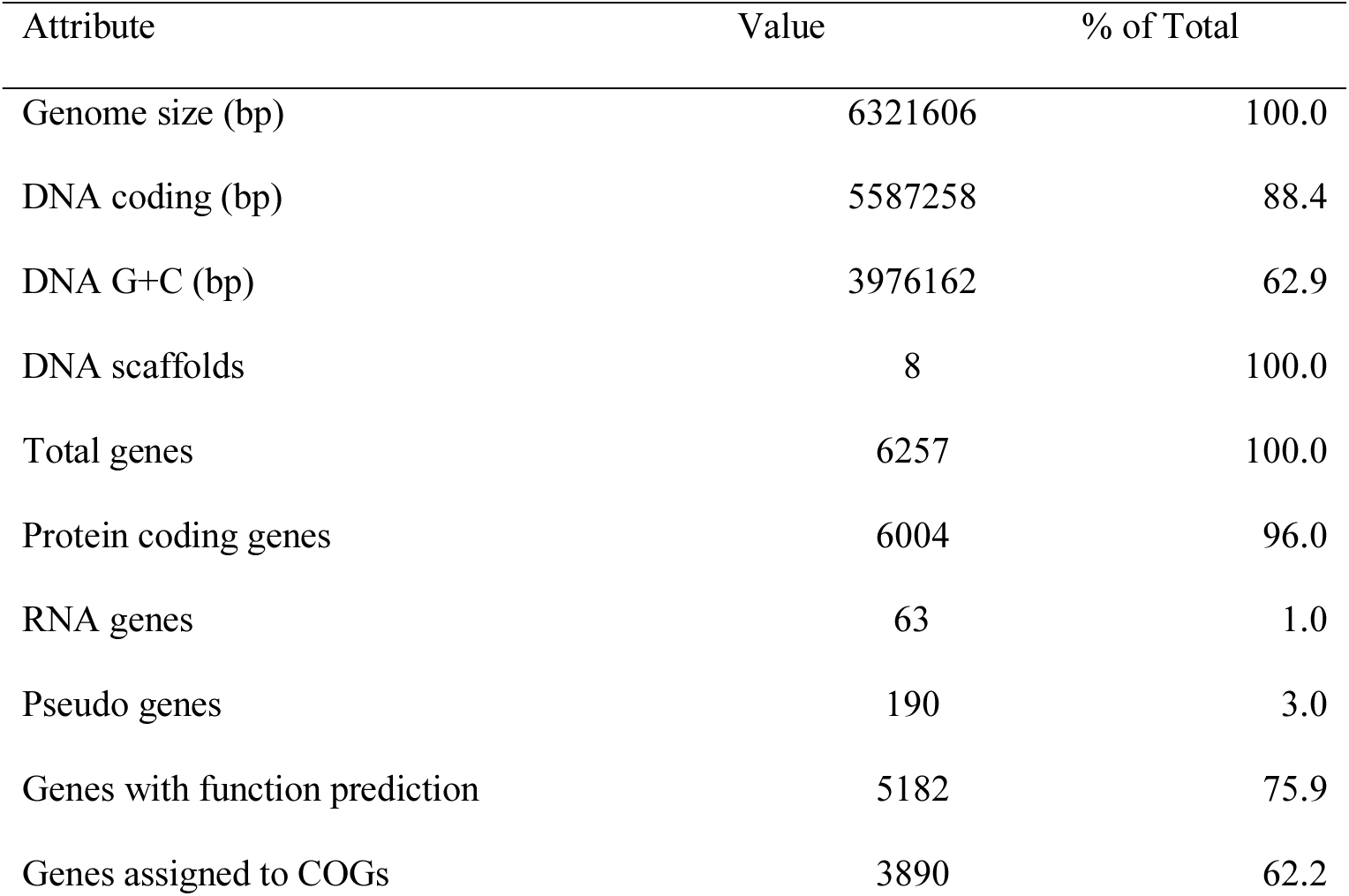
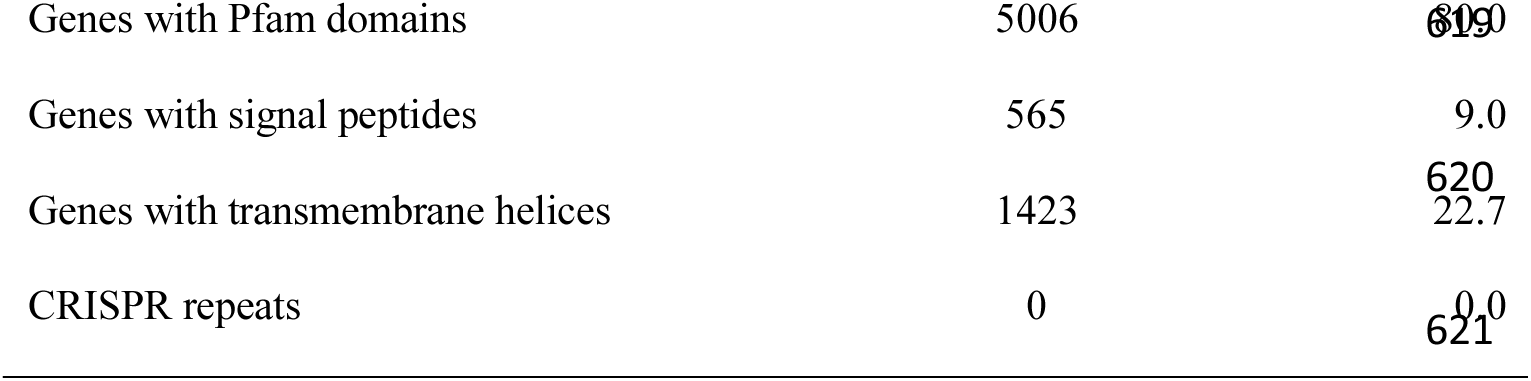
Genome statistics based on substrain MK1.

*Phylogenetic assignment of MSH1 to* Aminobacter niigataensis

A phylogenetic tree based on the 16S rRNA gene sequence indicating the position of MSH1 is shown in Figure 3. The 1,463 bp 16S rRNA gene sequence of MSH1 is 100% identical to that of *Aminobacter niigataensis* DSM 7050^T^ and 99.6-99.8% to those of other *Aminobacter* species. This is supported by whole-genome *in silico* digital DNA:DNA hybridization using TYGS, which reports that MSH1 (substrain DK1) is 82.5% (recommended *d4* formula) similar to *A. niigataensis* DSM 7050. (Supplementary Table S1). Finally, ANI values against all available *Aminobacter* genomes from NCBI (complete and incomplete assemblies; downloaded January 31, 2021), showed an ANI of 98% against *A. niigataensis* DSM 7050 (Figure 4). Based on these analyses, we reassign *Aminobacter* sp. MSH1 as *Aminobacter niigataensis* MSH1.

**Figure 3.**
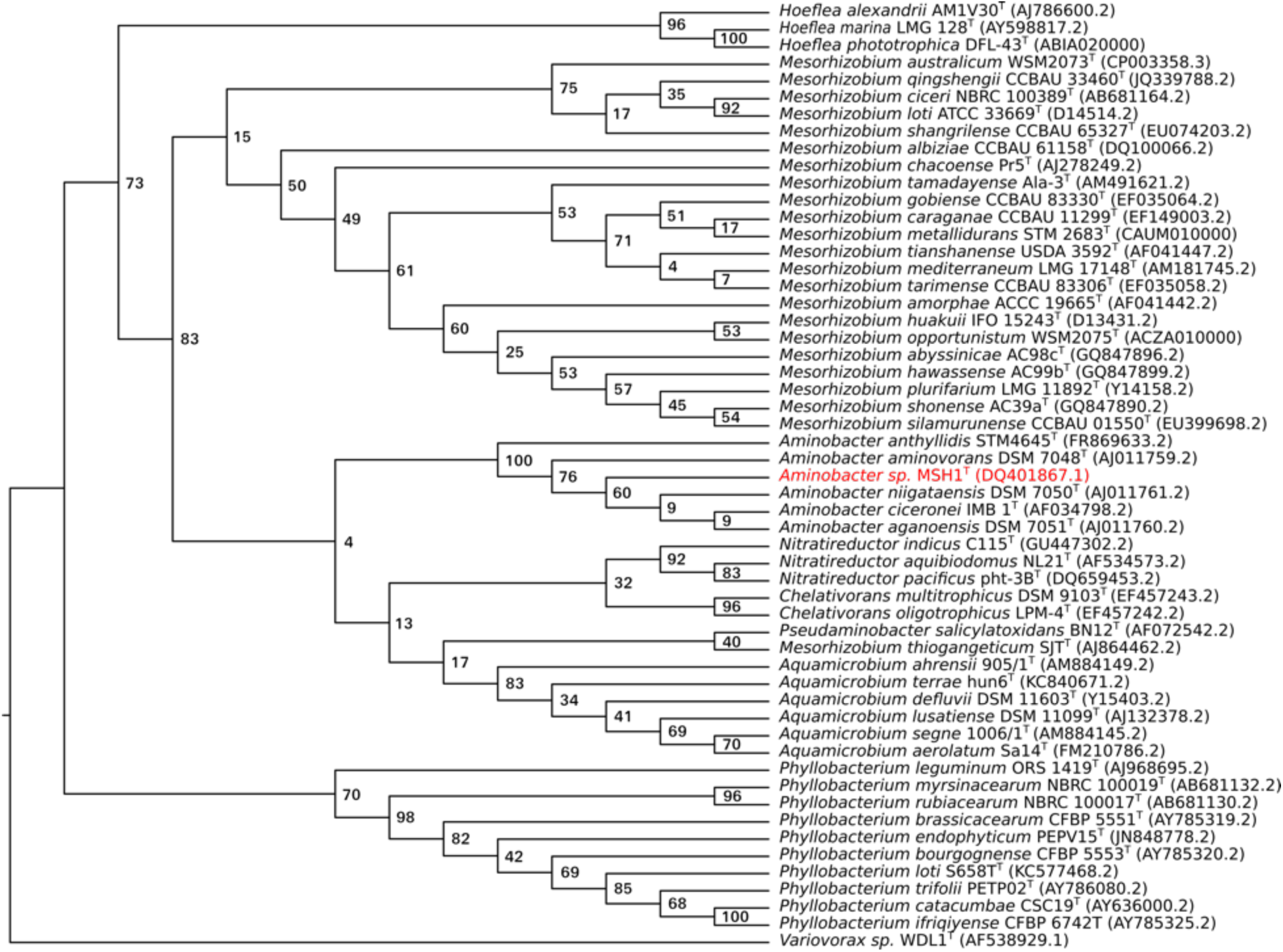
Phylogenetic relationships of *Aminobacter niigataensis* MSH1 based on the 16S rRNA gene sequence. Maximum likehood tree visualized as a cladogram with bootstrap values. This tree was created from a clustal-omega (30) multiple sequence alignment using 16S rRNA genes from the set of type strains available in the *Phyllobacteriaceae* family (NCBI accession numbers between parenthesis). The tree was inferred using PhyML (31) with a GTR substitution model and a calculation of branch support values (bootstrap value of 1000). The *Variovorax* sp. strain WDL1 was used as an outgroup (56).

**Figure 4.**
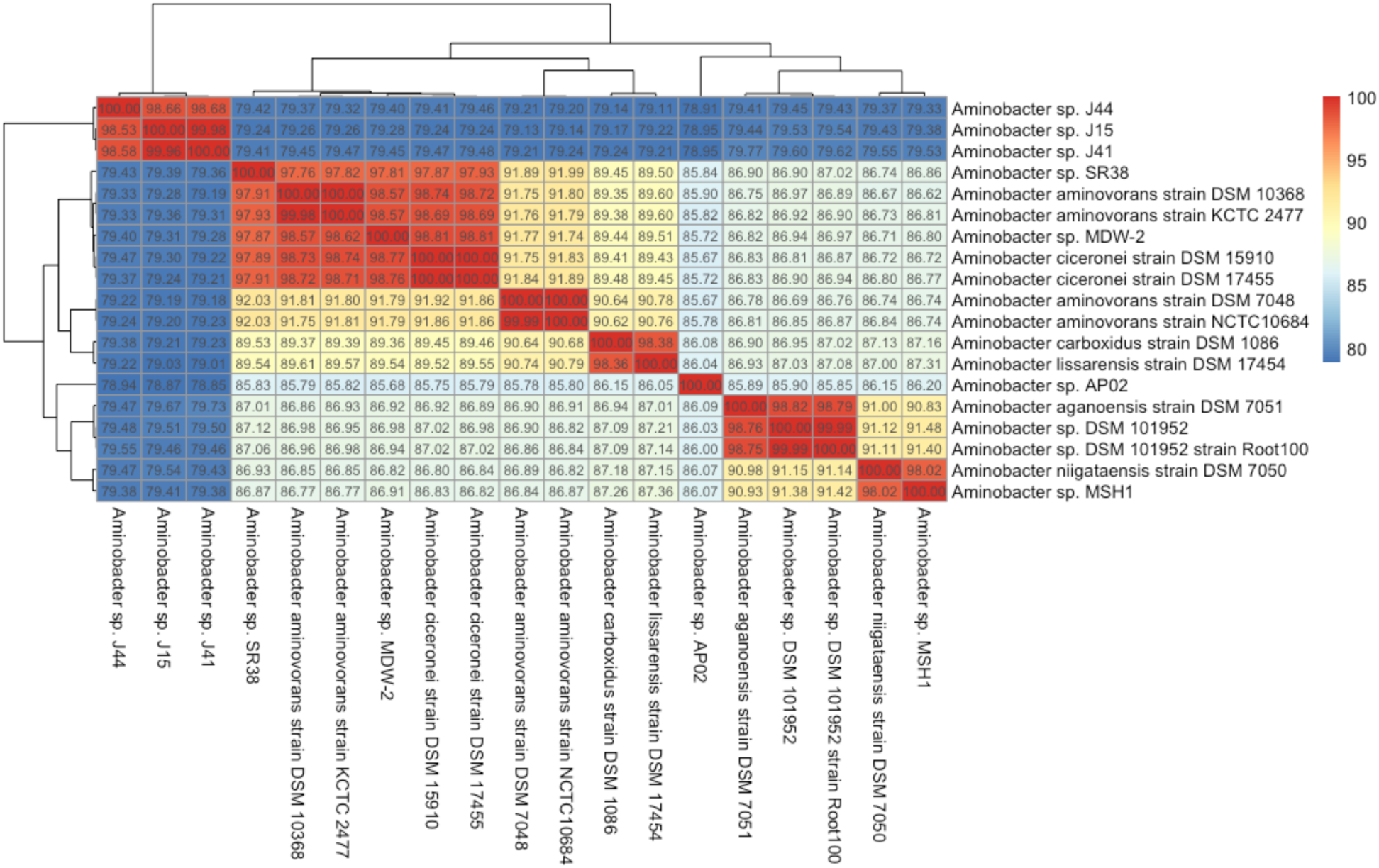
Heatmap of ANI values for all available *Aminobacter* genomes from NCBI (downloaded January 31, 2021). Genomes are clustered using hierarchical clustering of ANI values, as implemented in the R package “pheatmap” (v1.0.12).

### Chromosomally encoded metabolic features of MSH1

The chromosome of MSH1 possesses all genes required for glycolysis using the Embden-Meyerhof pathway and additionally possesses all genes for glucose metabolism through the Entner-Doudoroff pathway and the pentose phosphate pathway. It also contains all genes of the tricarboxylic acid cycle. MSH1 was previously shown to grow slower on succinate and acetic acid as carbon sources compared to glucose, fructose, and glycerol (43). MSH1 does not possess genes involved in carbon fixation which rules out autotrophic growth. MSH1 further displays the catechol *ortho*-cleavage pathway (44) and possesses genes for conversion of benzoate to catechol allowing the organism to grow on benzoate which was confirmed by culturing the strain on benzoate (data not shown). With regards to nitrogen metabolism, MSH1 contains a gene cluster that encodes the transmembrane ammonium channel AmtB as well as its cognate protein GlnK (45) for controlling ammonium influx in response to the intracellular nitrogen status, indicating that MSH1 can use mineral ammonia as a nitrogen source directly from its environment. In addition, MSH1 encodes for proteins involved in nitrate transport (NrtA and NrtT). The corresponding genes are located upstream of genes for assimilatory nitrate reduction (*nasDEA*) to ammonium suggesting that MSH1 can also use nitrate as a nitrogen source. Finally, ammonia is also released from amino acid metabolism and is further incorporated in L-glutamate for biosynthesis. Furthermore, MSH1 contains a gene cluster which combines a periplasmic dissimilatory nitrate reductase (*napAB),* the membrane-bound cytochrome *c* (*napC*) that is involved in electron transfer from the quinol pool in the cytoplasmic membrane to NapAB, *nirK* (nitrate reductase) and *norBC* (nitrix oxide reductase). However, *narG,* encoding the cytoplasmatic oriented dissimilatory nitrate reductase, is lacking. Dissimilatory nitrate reductases are associated with the cell membrane, and are typically involved in energy acquisition, detoxification, and redox regulation (46) NarG, located at the cytoplasmatic side of the cell membrane, is the typical respiratory nitrate reductase though its function can be replaced by NapAB in cases coupled to formate oxidation (46). However, this is unlikely since MSH1 is unable to grow under nitrate reducing conditions (6). In addition, *nosZ* for reduction of nitrous oxide to dinitrogen (47) is missing. The exact function of the gene cluster containing *napABC*, *nirK* and *norBC* is yet unknown. Besides direct uptake, for sulfur metabolism, MSH1 possesses two nearby located gene clusters encoding the ABC transporter complex CysUWA involved in sulfate/thiosulfate import. One of the two clusters appears directed to the uptake of sulfate while the other to thiosulfate uptake, since they respectively are linked with *sbp* and *cysP* (48). Both genes encode for the periplasmic protein that delivers sulfate or thiosulfate to the ABC transporter for high affinity uptake but Sbp binds sulfate and CysP thiosulfate. Furthermore, the chromosome contains all genes necessary for assimilatory sulfate reduction. The pathway reduces sulfate to sulfide involving ATP sulfurylase (CysND), adenosine 5’-phosphosulfate reductase (CysC), 3’-phosphoadenosine-5’-phosphosulfate reductase (CysH) and sulfite reductase (CisIJ). In addition, MSH1 contains *cysK* encoding O-acetylserine sulfhydrylase that incorporates sulfide into O-acetylserine to form cysteine (48). The assimilation of thiosulfate is less clear but MSH1 encodes for another homologue of CysK as well as several glutaredoxin proteins that are needed to incorporate thiosulfate in O-acetylserine and reductive cleavage reaction of its disulfide bond to form cysteine (48).

### Plasmids of MSH1

Besides the previously described IncP1-β and *repABC* plasmids, pBAM1 and pBAM2 (7, 19), substrain DK1 harbors the five pUSP1-5 plasmids (Figure 2), while substrain MK1 lacks pUSP2, pUSP3, pUSP4, and pUSP5. Catabolic genes on pBAM1 and pBAM2 enable MSH1 to mineralize the groundwater micropollutant BAM and use it as a source of carbon, nitrogen, and energy for growth. The amidase BbdA on pBAM1 transforms BAM to 2,6-dichlorobenzoic acid (DCBA) (7) which is further metabolized by a series of catabolic enzymes encoded by pBAM2 (19, 49). As previously discussed (19), the gene *bbdI* encoding the gluthatione dependent thiolytic dehalogenase responsible for removal of one of the chlorines from BAM together with *bbdJ* encoding gluthatione reductase, occur on pBAM2 in three consecutive, perfect repeats followed by a fourth, imperfect repeat. This, together with the placement of the BAM degradation genes on two separate plasmids (pBAM1 and pBAM2) and the bordering of the catabolic gene clusters by remnants of insertion sequences and integrase genes, suggests that the BAM catabolic genes in MSH1 have been acquired by horizonral gene transfer and then evolved to occur in their observed genomic organisation. In addition, pBAM2 has a considerably lower GC content of 56% compared to the chromosome and other plasmids which are between 60.0 and 64.4% (Table 2), which could indicate that pBAM2 was acquired from another, unknown, unrelated bacterium. It was previously shown that mineralization of DCBA is a common trait in bacteria in sand filters and soils, while BAM to DCBA conversion is the rate limiting step in BAM mineralization and is rare in microbial communities (50).

Like pBAM2, plasmids pUSP1, pUSP2, and pUSP3 belong to the *repABC* family. *repABC* replicons are known as typical genome components of *Alphaproteobacteria* species (51). The occurrence of more than one *repABC* replicon in one and the same genome has been described before and the plasmid family has been shown to exist of different incompatability groups. For instance, *Rhizobium etli* CFN42 has 6 *repABC* plasmids (52, 53).

Plasmids pBAM2, pUSP2, pUSP3, and pUSP4 contains Type IV secretion system (T4SS) genes (54), while pUSP1 does not. This indicates that pUSP1 is likely not self-transferable, unlike pBAM2, pUSP2, pUSP3, and pUSP4. Besides T4SS genes, plasmid pUSP4 contains a *mobABC* operon. The 31.6 kbp plasmid pUSP5 lacks conjugative transfer genes and appears to be a mobilizable plasmid with genes encoding a VirD4-like coupling protein and a TraA conjugative transfer relaxase likely involved in nicking at an *oriT* site and unwinding DNA before transfer. Furthermore, MOB-suite predicted that pBAM1, pUSP2, and pUSP4 have MOBP-type relaxase genes, while pUSP3 and pUSP5 have MOBQ-type relaxase genes. pBAM2 and pUSP1 were not predicted to have MOB-related genes.

### Specialized functions of plasmids pUSP1-5

In Table 3, all CDS of the different plasmids are categorized according to COGs. Half of the CDS annotated on plasmid pUSP1 (322 CDS) and pUSP2 (346 CDS) are genes primarily associated with the transport and metabolism of amino acids (20% and 12%, resp.), carbohydrates (6% and 6%, resp.) and inorganic compounds (10% and 3%, resp.), and genes for energy production and conversion (9% and 8%, resp.). For the plasmids pUSP3, pUSP4, and pUSP5, CDS categorized under the same COGs are lower than 18%. Together, pUSP1 and pUSP2 accounts for about 17% of all genes in MSH1 related to amino acid, carbohydrate transport and metabolism, and energy production and conversion in MSH1. The transport systems encoded by pUSP1 and pUSP2 include multiple ABC-transporters for N and/or S-containing organic compounds. For amino acids, carbohydrates and inorganic compound metabolism and transport, ABC-type transport systems are predicted for polar amino acids (arginine, glutamine), branched chain amino acids, and multiple sugars. In addition, transport systems for spermidine/putrescine, taurine, aliphatic sulphonates, dipeptides, beta-methyl galactoside, polysialic acid, and phosphate were predicted. Putative functions could be assigned by Prokka to 64.2%, 56.8%, 27.4%, 42.2%, and 34.3% of CDS for pUSP1, pUSP2, pUSP3, pUSP5 and pUSP5, respectively. On pUSP1, found in both MSH1 substrains, multiple genes could be assigned to metabolic subsystems by RAST. These include folate biosynthesis, cytochrome oxidases and reductases, degradation of aromatic compounds (homogentisate pathway), ammonia assimilation, and several genes related to amino acid metabolism. Some of these functions on pUSP1 do not have functional analogs on the chromosome, which may help to explain why pUSP1 was not lost in substrain MK1, but the other pUSP plasmids were. On pUSP2, which is absent in MK1 substrain, some genes are predicted to be involved in acetyl-CoA fermentation to butyrate, creatine degradation, metabolism of butanol, fatty acids, and nitrile, and a few miscellanoues functions. A large number of CDS on pUSP1 (19%), and pUSP2 (23%) are homologues to CDS on the chromosome and could be considered dispensable genes. However, although these CDS might be considered homologues, their functionality might differ considerably in terms of substrate specificity and kinetics.

**Table 3.** Percentage of genes associated with general COG functional categories in genome and replicons.

| Code | Description | Total | Chr | pBAM1 | pBAM2 | pUSP1 | pUSP2 | pUSP3 | pUSP4 | pUSP5 |
| --- | --- | --- | --- | --- | --- | --- | --- | --- | --- | --- |
| J | Translation, ribosomal structure and biogenesis | 2.9% | 3% | 0% | 0% | 2% | 1% | 0% | 0% | 0% |
| A | RNA processing and modification | 0.0% | 0% | 0% | 0% | 0% | 0% | 0% | 0% | 0% |
| K | Transcription | 7.3% | 7% | 5% | 9% | 10% | 7% | 9% | 5% | 18% |
| L | Replication, recombination and repair | 4.9% | 4% | 20% | 21% | 2% | 12% | 14% | 16% | 18% |
| B | Chromatin structure and dynamics | 0.1% | 0% | 0% | 0% | 0% | 0% | 0% | 0% | 0% |
| D | Cell cycle control, Cell division, chromosome partitioning | 0.7% | 1% | 2% | 2% | 2% | 1% | 2% | 3% | 6% |
| V | Defense mechanisms | 1.0% | 1% | 0% | 0% | 0% | 1% | 0% | 2% | 0% |
| T | Signal transduction mechanisms | 2.5% | 3% | 0% | 0% | 1% | 1% | 0% | 3% | 3% |
| M | Cell wall/membrane biogenesis | 3.8% | 4% | 0% | 0% | 2% | 1% | 2% | 2% | 0% |
| N | Cell motility | 0.6% | 1% | 0% | 0% | 0% | 0% | 0% | 0% | 0% |
| U | Intracellular trafficking and secretion | 2.5% | 2% | 24% | 17% | 0% | 3% | 11% | 22% | 3% |
| O | Posttranslational modification, protein turnover, chaperones | 3.0% | 3% | 0% | 17% | 1% | 0% | 0% | 0% | 0% |
| C | Energy production and conversion | 5.1% | 5% | 2% | 2% | 9% | 8% | 5% | 2% | 12% |
| G | Carbohydrate transport and metabolism | 4.1% | 4% | 0% | 6% | 6% | 6% | 2% | 2% | 0% |
| E | Amino acid transport and metabolism | 9.8% | 9% | 2% | 2% | 20% | 12% | 7% | 6% | 0% |
| F | Nucleotide transport and metabolism | 1.6% | 2% | 0% | 0% | 1% | 0% | 0% | 0% | 0% |
| H | Coenzyme transport and metabolism | 2.3% | 3% | 0% | 0% | 2% | 1% | 0% | 0% | 0% |
| I | Lipid transport and metabolism | 2.3% | 2% | 0% | 0% | 2% | 5% | 3% | 0% | 3% |
| P | Inorganic ion transport and metabolism | 5.8% | 6% | 0% | 0% | 10% | 3% | 0% | 0% | 3% |
| Q | Secondary metabolites biosynthesis, transport and catabolism | 1.9% | 2% | 5% | 4% | 5% | 6% | 1% | 0% | 0% |
| R | General function prediction only | 0.0% | 0% | 0% | 0% | 0% | 0% | 0% | 0% | 0% |
| S | Function unknown | 20.7% | 21% | 10% | 13% | 17% | 21% | 17% | 5% | 21% |
| - | Not in COGs | 16.9% | 17% | 29% | 8% | 9% | 12% | 27% | 33% | 15% |
| **CDS** |  | **6237** | **5277** | **41** | **53** | **322** | **346** | **100** | **63** | **34** |

Besides genes encoding conjugative transfer, plasmid replication, and plasmid stability functions, most genes on plasmids pUSP3, pUSP4, and pUSP5 could not be annotated with a function. However, several genes on pUSP3 may have functions related to metabolism of sugars, including inositol and mannose which were not tested in an earlier growth optimization experiment (43). On pUSP4, genes encoding a transmembrane amino acid transporter are situated next to an aspartate ammonia-lyase-encoding gene that enables conversion between aspartate and fumarate that may enter the tricarboxylic acid cycle, as described above. A cytochrome bd-type quinol oxidase, encoded by two subunit genes on pUSP5, also occurs in some nitrogen-fixing bacteria where it is responsible for removing oxygen in microaerobic conditions. Furthermore, a pseudoazurin type I blue copper electron-transfer protein is encoded by a gene on pUSP5, that may act as an electron donor in a denitrification pathway. A chromate transporter, ChrA, encoded by a gene on pUSP5 may confer resistance to chromate. Future studies should look into whether the lack of plasmids pUSP2-5 in substrain MK1 has phenotypic consequences, with regards to the predicted functions, including metabolism of sugars and aspartate, nitrogen metabolism, and resistance to chromate.

### Plasmid stability and chromosome polyploidy

The Illumina sequencing coverage of several plasmids relative to the chromosome (except for pBAM1) was lower than one, i.e., approximately 0.3 to 0.6 per chromosome. This suggests that either not all cells (only three to six out of ten) contain a copy of the same plasmid due to plasmid loss or that there are multiple copies of the chromosome. Previously, in the MK1 substrain, we observed that pBAM2 is not always perfectly inherited by the daughter cells in cultures grown in R2B and R2B containing BAM (15). To observe whether plasmid instability explained the copy number relative to the chromosome in the sequenced cultures, sequencing was performed directly on the cryo stock as well as on colonies directly derived from this, mimicking the sequenced cell preparation for whole genome sequencing. We hypothesized that if certain plasmids are not stably inherited (i.e. those with copy numbers 0.3 to 0.6), only part of the cell population will habour those plasmids and picking of multiple colonies from a plate will result in picking of some colonies that have lost one or more plasmids.

MSH1 was sequenced directly from the cryostock, from a single colony picked from R2A plates after spreading the cryostock, and from the broth R2B culture that had been inoculated with the same single colony from cryostock. Moreover, after spreading the latter R2B culture on an R2A plate, an additional 14 MSH1 colonies were picked for sequencing. Taking into account a plasmid coverage of 0.3 – 0.6 per chromosome, we expect that around half of the colonies would have lost one or more of the plasmids in the case of poor inheritance. However, only one of the colonies showed loss of a plasmid, i.e., plasmid pUSP1 (Figure 5) indicating polyploidy of the chromosome rather than unstable inheritance of plasmids. The loss of *repABC* megaplasmid pUSP1 shows that the possible metabolic features encoded by genes on pUSP1, as described above, are not essential for growth under these conditions, although, remarkably this is the only pUSP plasmid still present in substrain MK1. Interestingly, the plasmid/chromosome-ratio varied according to the growth medium from which DNA was isolated. When growing in R2B (broth), e.g. as done for DNA extraction for genome sequencing and from cryostock and R2B culture (first and third green rings, Figure 6), all plasmids, except pBAM1, have a copy number lower than one per chromosome. When DNA was extracted from colonies grown on R2A plates (though resuspended in PBS prior to DNA extraction), plasmid copy numbers were approx. one per chromosome, except for pBAM1 which has a copy number of approx. 2.5 per chromosome.

**Figure 5.**
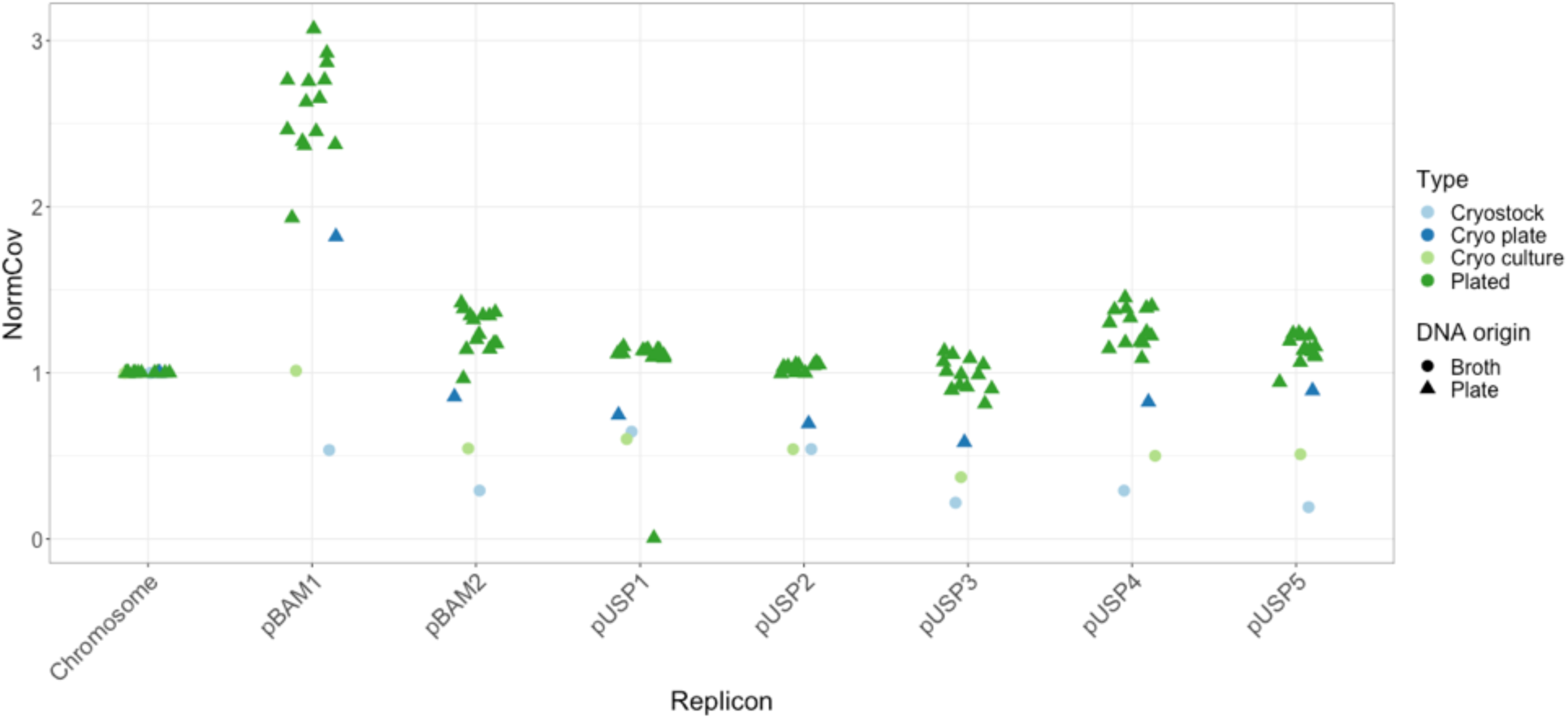
Coverage of replicons normalized to chromosome coverage (NormCov). A NormCov of 1 indicates a single copy per chromosome of a replicon. A NormCov above 1 indicates that there are more copies of a given plasmid than the chromosome per cell. Points have been slightly jittered horizontally to improve visualization of overlaps.

**Figure 6.**
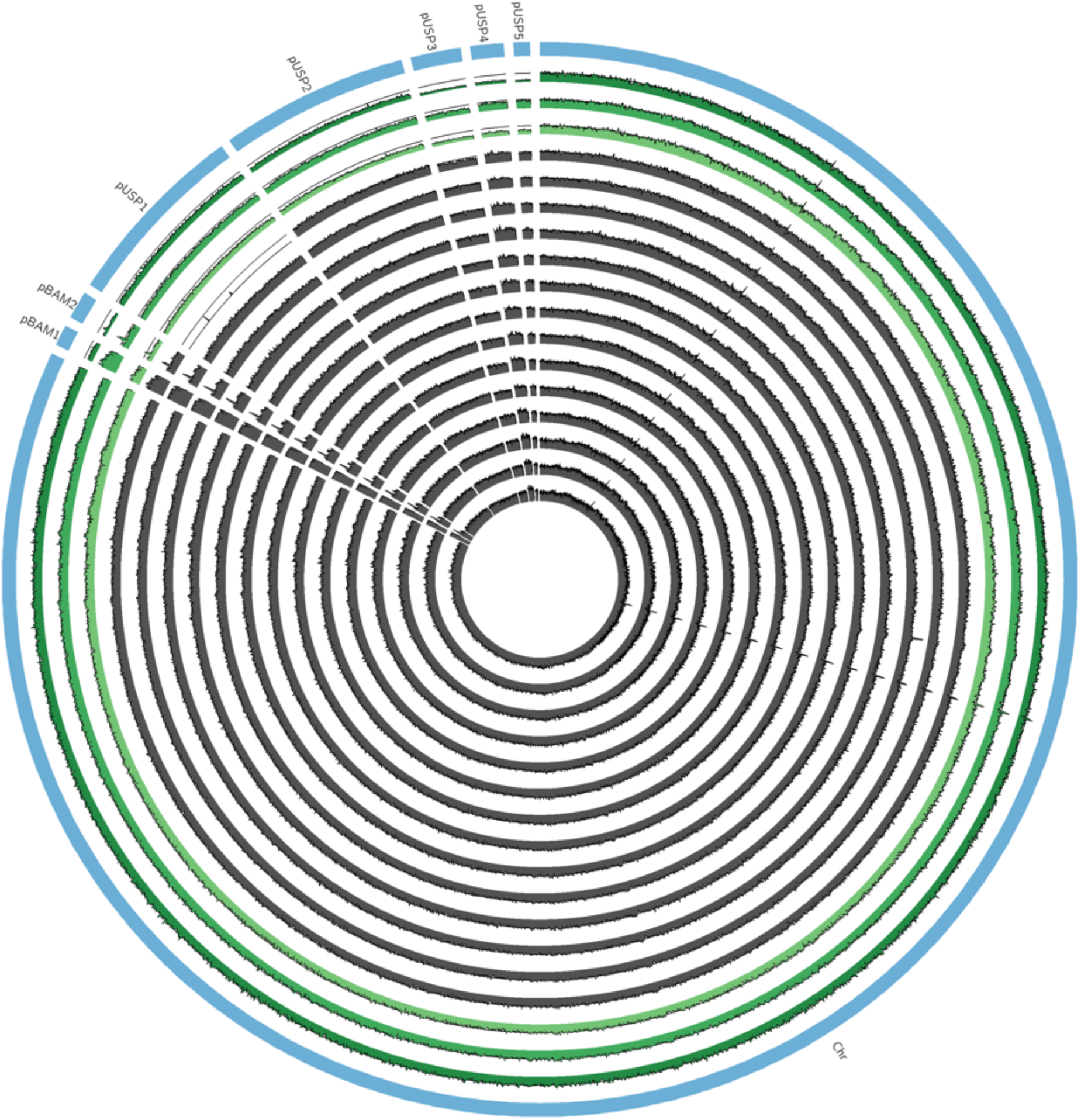
Illumina reads mapped to the chromosome and plasmids of MSH1 (DK1 substrain). The outer blue ring indicates the replicons. The inner rings show read mapping coverage of the replicons, normalized to the coverage of the chromosome per replicate, for all of the 17 sequenced replicates. The three first green rings show replicates cryostock, first colony from R2A plate, first colony from R2B broth, respectively. The subsequent 14 grey rings show the replicates that all originate from the same first colony. A solid line in the background of all tracks indicate the chromosome coverage line. Coverage above this line indicates a replicon copy number higher than 1 per chromosome and vice versa.

Except for the single loss of pUSP1, nothing here indicates unstable maintenance of plasmids and subsequent loss. Instead, our results indicate that MSH1 regulates the chromosome copy number according to whether it grows as planktonic bacteria or fixed on an agar plate. The results shown here can be explained by MSH1 being polyploid with regards to its chromosome when growing in broth media. Single-copy plasmids (e.g. pBAM2, pUSP1-5) will thereby have copy numbers lower than one relative to the chromosome, when growing in broth R2B. Polyploidy in prokaryotes have been described before, including in *Deinococcus*, *Borrelia*, *Azotobacter*, *Neisseria*, *Buchnera*, and *Desulfovibrio* (55) and may be quite overlooked in many other bacteria. *E. coli* in stationary phase was shown to have two chromosome copies after growing in rich, complex medium, but only 60% of the cells had two copies in stationary phase after slower growth in a synthetic medium (55). It was suggested that monoploidy is not typical for proteobacteria, and that many bacteria are polyploid when growing in exponential phase (55). Possible advantages offered by polyploidy include resistance to DNA damage and mutations, global regulation of gene expression by changing chromosome copy number, and finally polyploidy may enable heterozygosity in bacteria where genes mutate to cope with challenging condition while preserving a copy of the original genes. Despite the stability of the plasmids in MSH1, the MSH1 substrain MK1 lacks plasmids pUSP2-5 and a loss of pBAM2 was previously observed (15). Although pBAM2 encodes its own T4SS, the multiple loss of pBAM2 and pUSP2-5 in MK1 could be hypothetically explained by some uncharacterized plasmid codependence, where one loss leads to another. The dynamics of plasmid loss that has led to formation of substrain MK1 are still unknown.

## Conclusions

The full genome of *Aminobacter* sp. MSH1, re-identified here as *Aminobacter niigataensis* MSH1, consisting of a chromosome and seven plasmids, was determined combining both Nanopore and Illumina sequencing. Two smaller plasmids pBAM1 and pBAM2 were previously identified carrying the catabolic genes required for mineralization of the groundwater micropollutant BAM. Both the chromosome and the other five plasmids are described here for the first time. A plasmid stability experiment showed that most plasmids were stably maintained, with exception of a single loss event of plasmid pUSP1. Instead, the results indicate that MSH1 has a polyploid chromosome when growing in broth, thereby reducing plasmid copy numbers per chromosome to below one. When comparing the original strain MSH1 (DK1) and substrain MK1, we observed that plasmids pUSP2, pUSP3, pUSP4, and pUSP5 were below detection limits in MK1. Substrain MK1 may previously have lost these plasmids but maintained pUSP1, pBAM1, and pBAM2, thereby retaining its capacity to degrade BAM. Future studies on growth and degradation kinetics of the MSH1 and its substrain MK1 lacking several plasmids, can reveal if plasmids pUSP2-5 harbour unknown (favorable) functions or if they impose a metabolic burden on MSH1. This will help to elucidate which substrain is preferable for bioaugmentation.

## Funding

This project (LHH, LEJ, OH, and JAA) was funded by MEM2BIO (Innovation Fund Denmark, contract number 5157-00004B) at Aarhus University and DS by the KU Leuven C1 project no. C14/15/043 and the BELSPO IAP-project µ-manager no. P7/25 at KU Leuven. BH was supported by postdoctoral fellowship grant 12Q0218N and CL by an SB-FWO PhD fellowship 1S64718N. TKN was supported by AUFF NOVA project ORIGENE.

## Authors’ contributions

LHH, TKN, BH and DS initiated, planned, and funded the project. BH and TKN wrote the initial draft manuscript. BH, TKN, and CL performed genome sequencing, assembly annotation, and data analyses. OH performed the plasmid stability experiment. VvN, RL, DS, LEJ, JA, and LHH critically reviewed the paper, assisted in data interpretation, and coordinated experiments.

---

**Supplementary Table S1.**
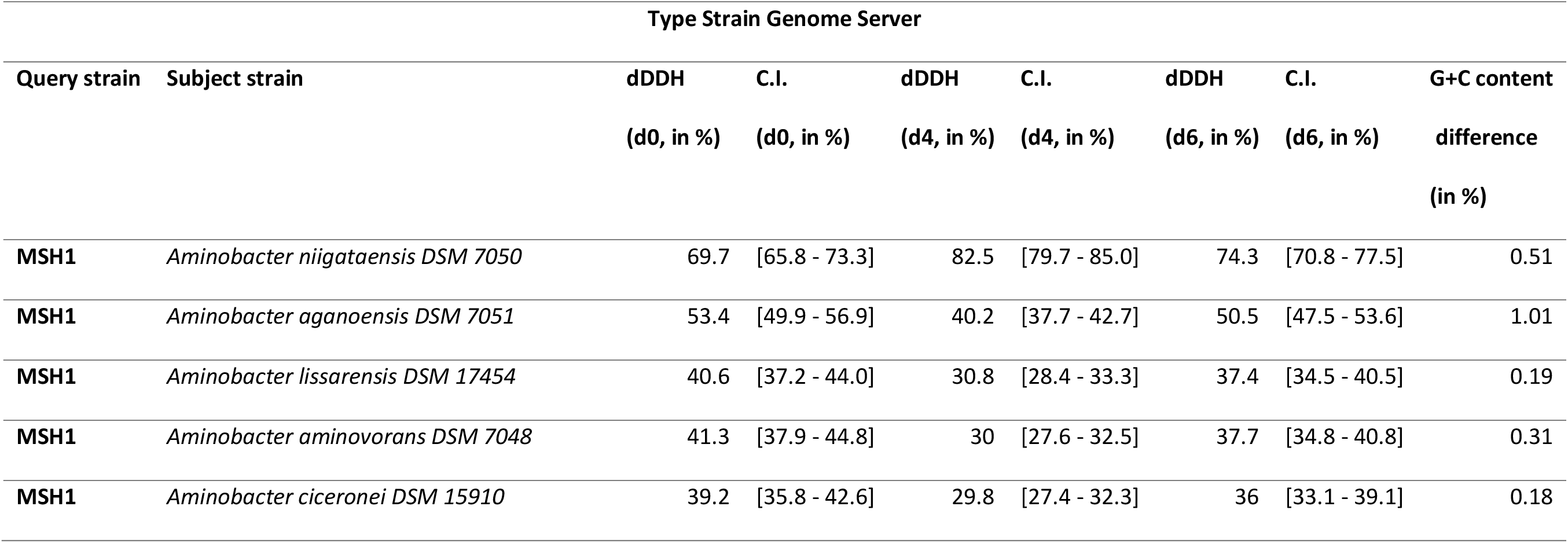
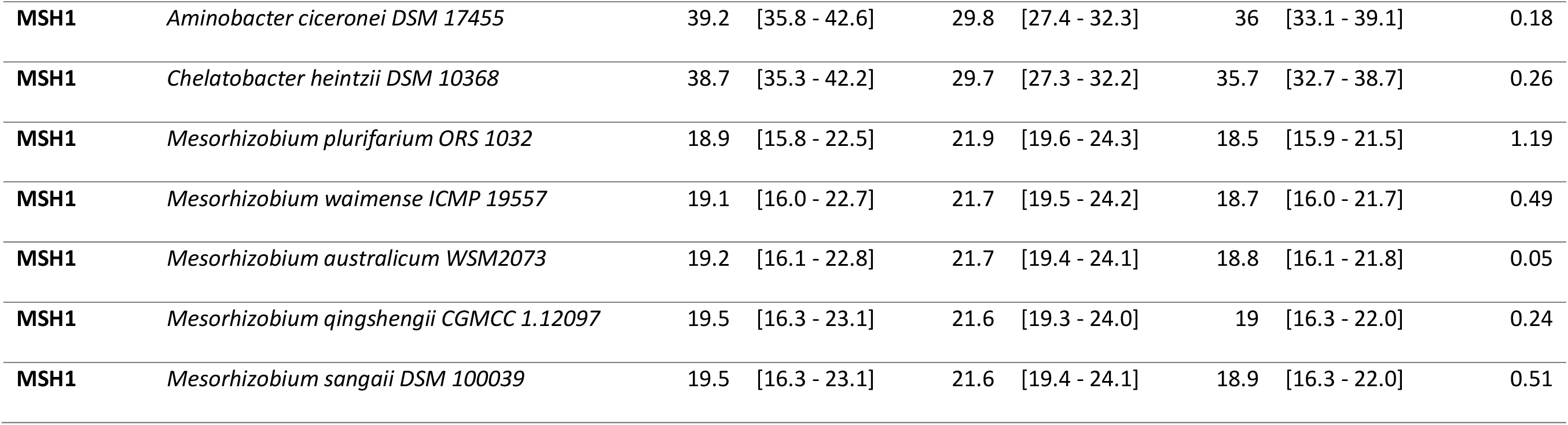
Digital DNA:DNA hybridization (dDDH) results from the online Type Strain Genome Server (TYGS) analysis. d0, d4, d6 refers to different algorithms used TYGS. Formula d0 (a.k.a. GGDC formula 1): length of all HSPs divided by total genome length. Formula d4 (a.k.a. GGDC formula 2): sum of all identities found in HSPs divided by overall HSP length. Formula d6 (a.k.a. GGDC formula 3): sum of all identities found in HSPs divided by total genome length. C.I.: Confidence interval.

